# The sterol transporter STARD3 transports sphingosine at ER-lysosome contact sites

**DOI:** 10.1101/2023.09.18.557036

**Authors:** Pia Hempelmann, Fabio Lolicato, Andrea Graziadei, Ryan D. R. Brown, Sarah Spiegel, Juri Rappsilber, Walter Nickel, Doris Höglinger, Denisa Jamecna

## Abstract

Sphingolipids are important structural components of membranes. Additionally, simple sphingolipids such as sphingosine are highly bioactive and participate in complex subcellular signaling. Sphingolipid deregulation is associated with many severe diseases including diabetes, Parkinson’s and cancer. Here, we focus on how sphingosine, generated from sphingolipid catabolism in late endosomes/lysosomes, is reintegrated into the biosynthetic machinery at the endoplasmic reticulum (ER). We characterized the sterol transporter STARD3 as a sphingosine transporter acting at lysosome-ER contact sites. Experiments featuring crosslinkable sphingosine probes, supported by unbiased molecular dynamics simulations, exposed how sphingosine binds to the lipid-binding domain of STARD3. Following the metabolic fate of pre-localized lysosomal sphingosine showed the importance of STARD3 and its actions at contact sites for the integration of sphingosine into ceramide in a cellular context. Our findings provide the first example of inter-organellar sphingosine transfer and pave the way for a better understanding of sphingolipid – sterol co-regulation.

## Introduction

Lysosomes are bilayered organelles that facilitate cellular catabolic reactions and play essential roles in cellular signaling. ^1^ Defects in lysosomal function were linked to various diseases such as neurodegenerative disorders, cancer, inflammation and lysosomal storage diseases, many of which feature a strong lipid accumulation phenotype. ^2–6^

In this light, lysosomal lipid recycling is of significant interest but has so far only been elucidated for sterols, mainly because more tools and methods exist to follow sterol movement in the cell.^7^ Endocytosed dietary cholesterol is released from low density lipoprotein (LDL) particles and incorporated into the lysosomal limiting membrane via transmembrane transporters such as Niemann-Pick C1 (NPC1) and Lysosome membrane protein 2 (LIMP-2/SCARB2) for further export. ^8–10^ Membrane contact sites (MCS) between lysosomes and the ER participate in the export of cholesterol into the ER for its subsequent trafficking throughout other organelles. ^8,11,12^ Such contact sites are also needed to supply lysosomes with ER-derived cholesterol. ^13–15^

In contrast, the lysosomal export of sphingolipids is much more enigmatic. Sphingolipids are particularly enriched at the plasma membrane and undergo constitutive turnover throughout the cell. ^16,17^ Catabolism of complex sphingolipids in lysosomes produces sphingosine, which is then exported by unknown mechanisms to the ER and subsequently to the Golgi for the synthesis of new sphingolipids. ^16,18^

In our previous work, we introduced chemical biology tools to manipulate and follow cellular sphingosine using lipid probes with photocrosslinkable and clickable (pac) features. ^19,20^ The photocrosslinkable functionality enables the creation of stable protein-lipid complexes by irradiation with UV light, whereas the clickable functionality enables click chemistry reactions with various tags for visualization and further utilization in downstream applications such as mass spectrometry. We subsequently equipped pac-lipid probes with organelle targeting groups that allowed resolving cellular lipid transport and metabolism with high spatial and temporal precision. Using lysosome targeted pac-sphingosine (lyso-pacSph), we recently identified sterol transporters NPC1 and, to a lesser extent, LIMP-2 as involved in sphingosine efflux from the lysosome. ^21^ Both proteins feature intramolecular tunnel domains that channel cholesterol and sphingosine into the lysosomal limiting membrane. ^9,10^ From there, cholesterol trafficking proceeds further into the target membranes, such as the ER. This step is facilitated by dedicated transporters such as Oxysterol-binding protein (OSBP) and Oxysterol-binding protein-related protein 1(ORP1L). ^8,11–15^ For sphingosine, the proteins and mechanisms involved in its further export are still elusive.

In our previous study ^19^, a mass spectrometric screen for putative sphingosine interactors identified StAR-related lipid transfer protein 3 (STARD3). STARD3 is a 4 transmembrane domain-containing protein residing in the limiting membrane of late endosomes and lysosomes, initially discovered as highly overexpressed in metastatic cancer. ^22^ STARD3 forms organelle contacts with the ER by interacting with Vesicle-associated membrane protein-associated (VAP) and Motile sperm domain-containing (MOSPD) proteins ^23,24^ and was shown to transport cholesterol from the ER to lysosomes via its StART-domain. ^13^

In this study, we utilize pac-sphingosine and its organelle targeted version lyso-pacSph in cellular and *in vitro* contexts alongside lipidomics analyses and molecular dynamics simulations to show that STARD3 can traffic sphingosine for its reuse in the sphingolipid biosynthetic pathway.

## Results and Discussion

### STARD3 binds sphingosine inside its lipid binding cavity

As a first step, we confirmed the binding of Sph to STARD3 by using pac-lipids, as illustrated in Fig. 1A. Immunoprecipitation of crosslinked, stably overexpressed STARD3 in HeLa WT cell lysates showed a clear interaction with pacSph, but not with the control lipid pac-fatty acid (pacFA, Fig. 1B, quantified in Fig. 1C). Therefore, we rule out unspecific interaction of STARD3 with the single-chain lipids in the membrane. To control for the technical background of the streptavidin-mediated immunoprecipitation, we employed non-crosslinked controls (-UV), in which STARD3 was not pulled down. To further verify that the detected interaction was not a byproduct induced by the preparation of cell lysates, we employed a lysosomal-targeted variant of the pacSph probe (lyso-pacSph ^21^) in intact cells. Here, the photocrosslinking step was performed on living HeLa SGPL1^-/-^ cells ^25^ to avoid the breakdown of the released pacSph by sphingosine1-phosphate lyase (SGPL1). We could again detect a successful crosslinking between pacSph and STARD3, which is even increased in conditions where ceramide synthases are inhibited using the ceramide synthase inhibitor Fumonisin B1 (FB1, Supplementary Fig. 1). Altogether, these crosslinking immunoprecipitation experiments point towards an interaction between STARD3 and sphingosine, rather than its metabolites.

**Fig. 1:**
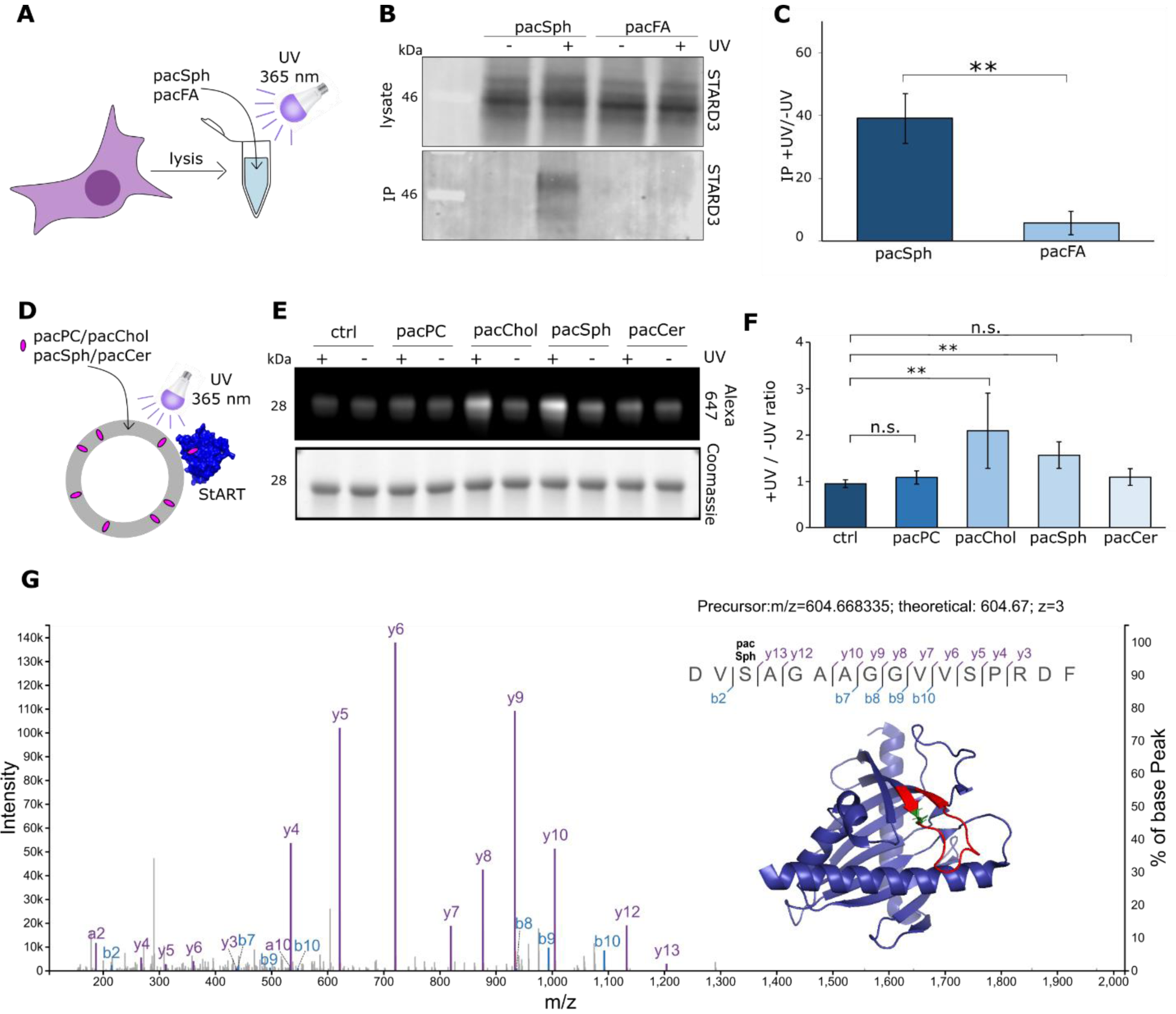
Sphingosine binds to STARD3 **A** Schematic illustration of pacSph/pacFA crosslinking experiments. **B** Immunoprecipitation of STARD3 and pacSph/pacFA in cell lysates. Cellular lysates of cells expressing FLAG-STARD3 were incubated with 10 µM pacSph or pacFA for 1 h at 37 °C and crosslinked with UV light (365 nm). Crosslinked protein-lipid complexes were conjugated with biotin-azide and bound to streptavidin-coupled beads. Eluted protein-lipid complexes were analyzed by western blotting using a STARD3-specific antibody. **C**. Quantification of +UV/-UV ratios of five independent experiments described in B. Welch two sample t-test was performed with **p < 0.01. **D** Schematic illustration of *in vitro* crosslinking experiments. **E** Crosslinking profile of purified StART domain of STARD3. Liposomes containing 1.5 mol% of pac-lipids were incubated with 1 µM STARD3-StART and UV-crosslinked. Protein-lipid complexes were subjected to click reaction with AF647-picolyl azide and visualized by SDS-PAGE. In-gel fluorescence of AF647 was normalized to Coomassie staining of loaded protein. +UV/-UV signal ratio was quantified in **F** and Welch two sample t-tests were performed with n.s. p > 0.05 and **p < 0.01 **G** Fragmentation mass spectrum identifying S334 in STARD3 covalently bound to pacSph (monoisotopic modification mass 307.2513). Spectral annotation tolerance: 5ppm. b,y,a fragment matches are annotated.

To reconstitute this binding *in vitro*, we purified the lipid-transporting StART domain of STARD3 and used pac-lipids incorporated in liposomes. Liposomes and purified StART domain were UV-crosslinked as depicted in Fig. 1D. Non-crosslinked samples (-UV) were used as a negative control. The click reaction with the fluororescent AF647-picolyl-azide allowed us to measure in-gel fluorescence of the crosslinked complexes on SDS-PAGE (Fig. 1E). Calculating the fluorescence ratio between crosslinked samples and -UV background, we established the interaction profile of the StART domain with a variety of commercially available pac-lipids such as phosphatidylcholine (pacPC), cholesterol (pacChol), sphingosine (pacSph) and ceramide (pacCer), as shown in Fig. 1F. As expected, we found that STARD3-StART crosslinked well with pacChol. In addition, we observed a similarly strong interaction with pacSph, highlighting the dual specificity of STARD3 for both cholesterol and sphingosine-derived probes. No significant interaction was detected with pacPC or pacCer, indicating that the crosslinks observed in cells are specifically attributed to the interaction with the StART domain rather than interactions involving the transmembrane domain.

Having obtained stable crosslinks between the purified domain and pacSph, our next objective was to determine the specific binding site of pacSph within the domain. To achieve this, we employed crosslinking mass spectrometry (MS) on the protein-lipid complexes. The crosslinking MS spectra unveiled a pacSph-modified peptide located near the entrance of the lipid transport cavity (Fig. 1G, modified peptide highlighted in red). Further computational analysis of the MS results showed that pacSph conjugated to serine at position 334 (shown in green sticks). Altogether, our data show that STARD3 can bind sphingosine within its lipid-binding StART domain, which suggests that the observed crosslinks stem from trafficking rather than regulatory interactions of Sph with STARD3.

### STARD3 delivers lysosomal sphingosine for reuse in sphingolipid biosynthesis

To investigate whether STARD3 can transfer sphingosine in a cellular context, we again employed the lyso-pacSph probe. As this probe is initially localized in lysosomes in its inactive form, releasing it by a flash of light gives a well-defined starting point for pulse-chase experiments, in which all downstream metabolites can be visualized on thin-layer chromatography (TLC)^20^. Lysosomal Sph can either be used in the sphingolipid recycling pathway, where it is incorporated into ceramide species at the ER followed by SM biosynthesis at the Golgi, or it is degraded through phosphorylation to sphingosine-1-phosphate (S1P) and cleavage by SGPL1, as illustrated in Fig. 2A. The breakdown product of SGPL1 action, hexadecanal, can be oxidized and introduced into abundant glycerolipids such as PC. In cells stably overexpressing (OE) STARD3, as well as STARD3 knock-out (KO) cells, the relative abundance of all lyso-pacSph derived metabolites was analyzed by TLC (Fig. 2B). Quantification of the band intensities (Fig 2C) showed a prominent increase of SM, representing the incorporation of lysosomal Sph into sphingolipid biosynthesis. In contrast, PC levels, indicative of sphingosine entering the SGPL1 breakdown pathway, significantly decreased in STARD3 OE conditions.

**Fig. 2:**
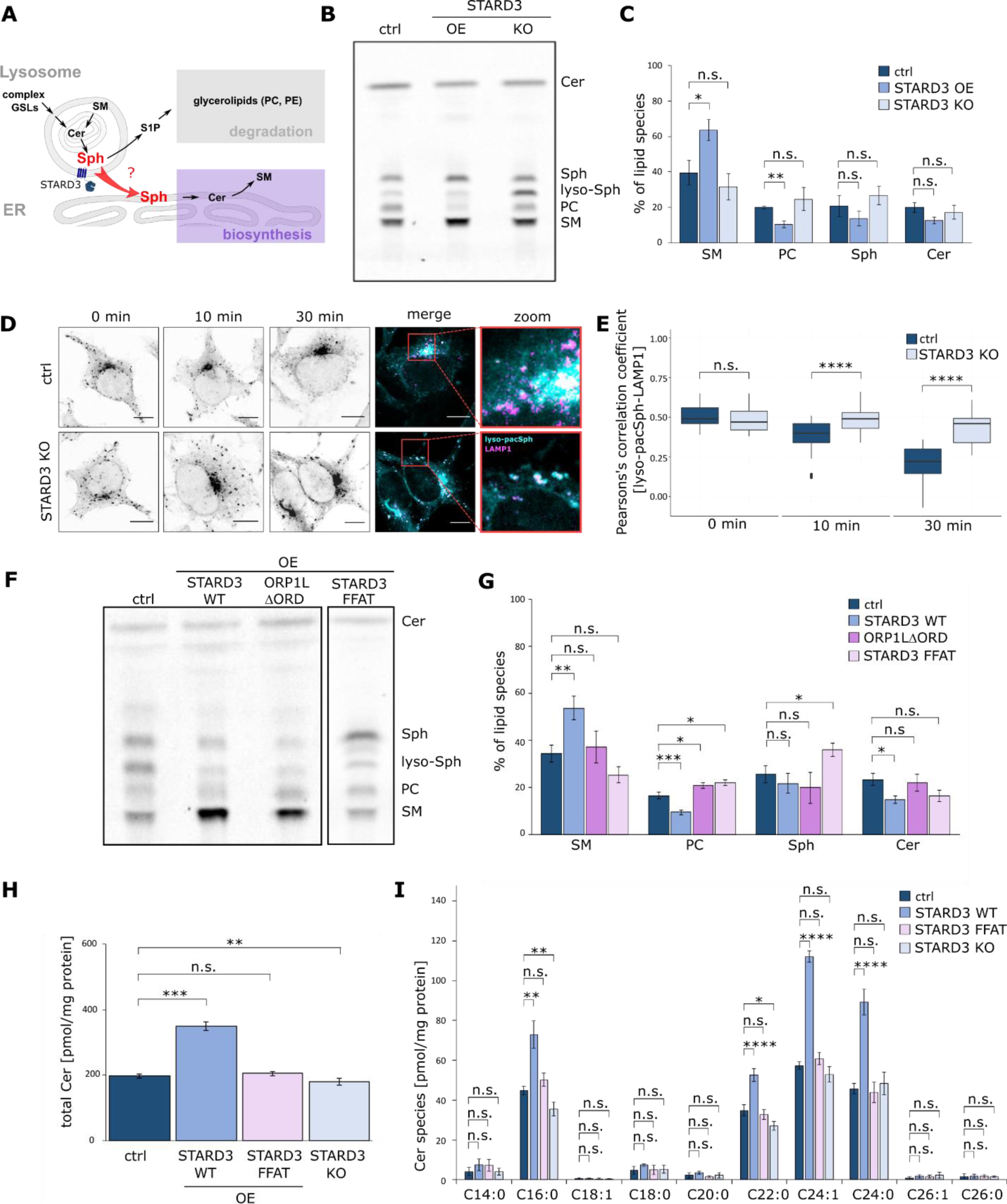
STARD3 delivers Sph for utilization in the biosynthetic pathway at the ER. **A** Schematic illustration of subcellular Sph metabolism. **B** Thin-layer chromatography of lyso-pacSph in ctrl, STARD3 OE and KO cells. Cells labeled with lyso-pacSph (5 µM, 17 h) were uncaged for 90s and chased for 30 min. Extracted lipids were visualized by coumarin azide and separated by TLC. Different lipid species were identified using lipid standards and quantified in **C** by calculating the ratio of a certain lipid species to the total lipid signal. Welch two sample t-tests were performed with n.s. p > 0.05, *p < 0.05 and **p < 0.01. **D** Confocal microscopy images of ctrl and STARD3 KO cells labeled with 10 µM lyso-pacSph. Scale bars, 10 µm. Lyso-pacSph was uncaged by UV light (405 nm) and crosslinked (365 nm) to proteins in close proximity 0 - 30 min post uncaging. Lyso-pacSph metabolites were conjugated to an Alexa488 fluorophore, and lysosomes were visualized using an immunostaining against LAMP1. Single microscopy images represent subcellular lyso-pacSph distribution, while merge images show lyso-pacSph (cyan) and LAMP1 (magenta) co-localization. **E.** Pearson’s correlation coefficient of lyso-pacSph and LAMP1 co-localization. Welch two sample t-tests were performed between ctrl and STARD3 KO conditions with n.s. p > 0.05 and ****p < 0.0001. **F** TLC analysis of lyso-pacSph in ctrl, STARD3 OE, ORP1LΔORD OE, and STARD3 FFAT mutant OE cells. TLC assay and lipid species quantification, shown in **G**, was performed as described above. Welch two sample t-tests were performed with n.s. p > 0.05, *p < 0.05, **p < 0.01 and ***p < 0.001. Sphingolipid mass spectrometry of **H** total Cer levels and **I** Cer species was performed using total cell lipid extracts of ctrl, STARD3 OE, STARD3 FFAT mutant OE and STARD3 KO cells. Welch two sample t-tests were performed in H and one-way ANOVA with Tukey post hoc tests were performed in I with n.s. p > 0.05, **p < 0.01 and ***p < 0.001.

On the other hand, STARD3 KO resulted in a slight, but not significant increase in PC and Sph levels (Fig. 2C), potentially pointing at a defect in utilization of sphingosine for sphingolipid biosynthesis. To investigate the impact of STARD3 KO on lyso-pacSph with a different method, we looked at the subcellular localization of the probe in fluorescence microscopy experiments. To this end, we incubated control and STARD3 KO cells with lyso-pacSph. We traced the localization of the probe 0, 10, and 30 min after uncaging through the addition of a fluorophore by click chemistry (Fig. 2D). While the initial lysosomal localization of the uncaged sphingosine probe changed to internal membrane and Golgi staining after 30 min post uncaging in control cells, STARD3 KO showed prolonged lysosomal staining of the liberated sphingosine probe. Immunofluorescence staining for the lysosomal marker protein LAMP1 (for co-localization, see Supplementary Fig. S2) allowed us to quantify the lysosomal export of lyso-pacSph over time by calculating the Pearson’s correlation coefficient of lyso-pacSph and LAMP1 (Fig. 2E). Indeed, we found an increased colocalization of lyso-pacSph and LAMP1 after 10 and 30 min in STARD3 KO conditions, pointing towards delayed export of lysosomal Sph when STARD3 is not available.

Having shown that overexpression of STARD3 increases the reutilization of Sph in the biosynthetic pathway of SM and that its absence leads to Sph accumulation in lysosomes, we next investigated whether these effects stem from the lipid transporting activity of STARD3 or from its function as a tether between ER and lysosomes. ^13^ Therefore, we employed a truncated version of another lysosome-ER tether, ORP1L (ORP1LΔORD), which was shown to increase contacts between lysosomes and ER without having lipid transport activity itself. ^26^ In cells overexpressing such ORP1LΔORD, we could not observe the increased incorporation of Sph into SM or the decreased flux towards PC as observed for STARD3 overexpression (Fig. 2F, quantified in G). We conclude that it is not the tethering ability of STARD3 that is responsible for the increased transfer of the lysosomal Sph backbone towards SM biosynthesis, but rather its actions as a Sph transporter.

Conversely, we investigated whether tethering of STARD3 to the ER is required for Sph transport by employing a STARD3 mutant with a defective FFAT motif, thus unable to bind to ER-resident VAP or MOSPD2 proteins. ^23^ Overexpressing such mutant and performing the lyso-pacSph TLC assay again showed no increased incorporation of Sph into SM or decreased incorporation into PC (Fig. 2F, quantified in G), highlighting the necessity of functional contacts for sphingosine transfer.

Having investigated the acute effects of STARD3 overexpression on the trafficking of the lyso-pacSph probe, we next investigated the effects of STARD3 on the endogenous sphingolipidome. Mass spectrometry analysis of control, STARD3-deficient cells as well as cells stably overexpressing a WT or FFAT-mutant variant of STARD3 (Fig. 2H) showed that the levels of the direct metabolite of Sph, ceramide, were elevated in STARD3-overexpressing cells and decreased in STARD3-deficient cells. The FFAT mutant did not influence ceramide levels, whereas STARD3 KO conditions resulted in slight, but significantly lower ceramide levels. Levels of endogenous SM species (Supplementary Fig. S3A and B) were not impacted by either overexpression or knock-out of STARD3. A closer investigation of ceramide species of different fatty acid chain lengths showed that the major long-chain ceramides C24:0 and C24:1, produced by the actions of CerS2, are particularly affected by STARD3 overexpression. This likely reflects the high expression levels of CerS2 in this cell line. ^27^

Altogether, our data are consistent with the hypothesis that STARD3 traffics Sph from lysosomes to the ER for use in the sphingolipid biosynthetic pathway, of which ceramide is the first step.

### Sphingosine and cholesterol trafficking functions of STARD3 compete with each other

Having now established STARD3 as a sphingosine transporter, we were curious about the interplay with its previously established function in cholesterol trafficking. STARD3 was shown to transfer cholesterol inside its StART domain, and a cavity-deficient mutant M307R/N311D (MR/ND) was characterized to no longer traffic sterols. ^13^ To investigate whether the sphingosine trafficking actions of STARD3 were similarly affected, we created cells stably overexpressing the MR/ND mutant as well as STARD3 lacking the entire StART domain (ΔStART). Again, we labeled these cells with lysosome-prelocalized sphingosine and analyzed the resulting metabolites by TLC following a 30 min chase upon uncaging (Fig 3A, B). In contrast to WT STARD3, overexpressing MR/ND and ΔStART mutants did not promote the synthesis of more complex pac-lipids, and the metabolic profile of lyso-pacSph after 30 min chase was not different from control cells. This STARD3-mediated sphingosine transfer deficiency in both conditions indicates that the cholesterol and sphingosine transport functions of STARD3 depend on the same structural features.

**Fig. 3.**
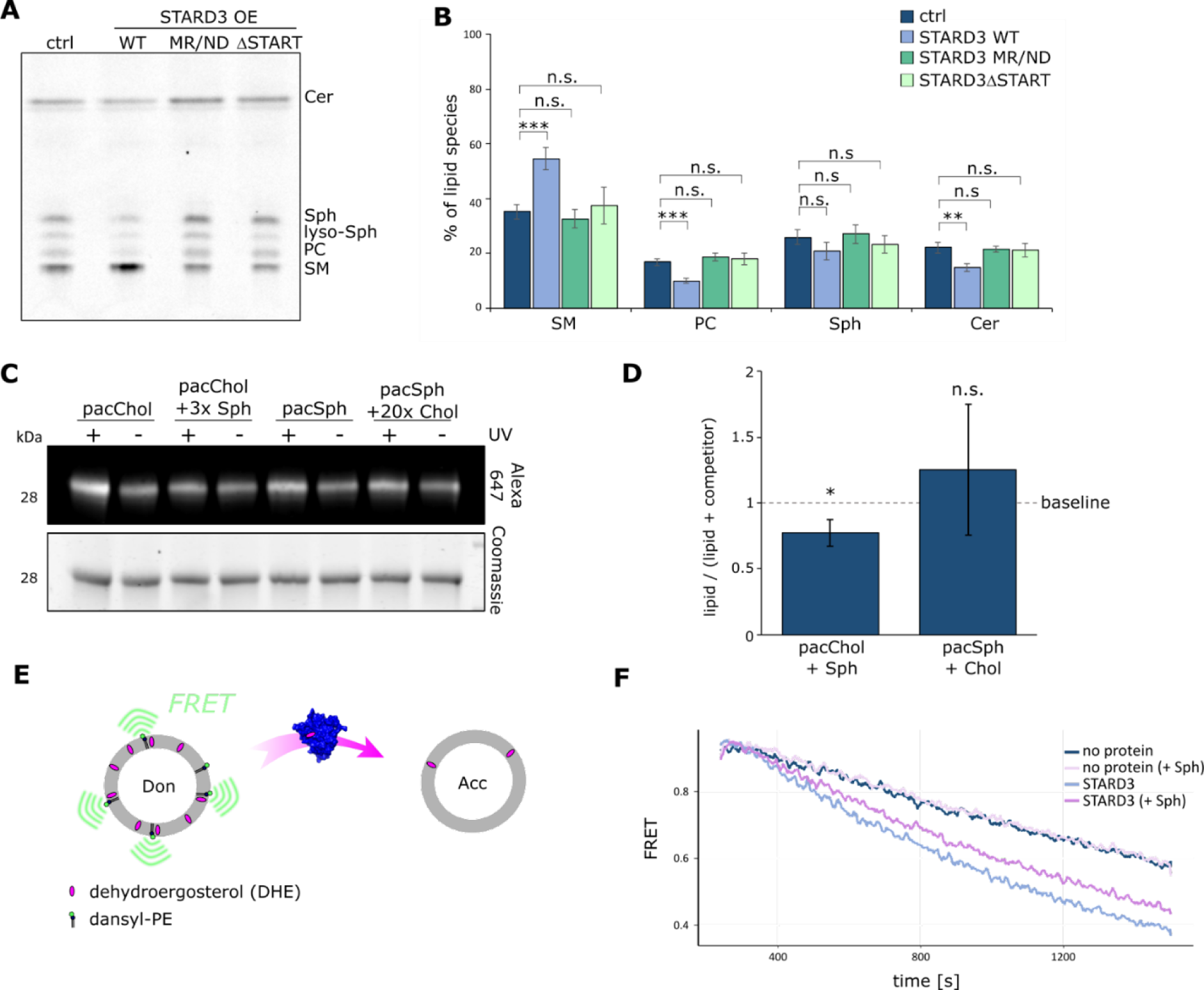
Sph and sterol transfer of STARD3 are in competition. **A** Thin-layer chromatography of lyso-pacSph in ctrl, STARD3 OE cells (WT, MR/ND mutant, and ΔStART). Cells labeled with lyso-pacSph (5 µM, 17 h) were uncaged for 90 sec by UV light (405 nm) and chased for 30 min. Extracted lyso-pacSph metabolites were conjugated to coumarin-azide and separated by TLC. Different lipid species were identified using lipid standards and quantified in **B** by calculating the ratio of a certain lipid species to the total lipid signal. Welch two sample t-tests were performed with n.s. p > 0.05, **p < 0.01 and ***p < 0.001. **C** Crosslinking competition profile of StART domain with pacChol/pacSph and natural sphingosine/cholesterol. Liposomes with 1.5 mol% pac-lipid and an excess of natural lipid (3-fold excess of sphingosine or 20-fold excess of cholesterol, respectively) were incubated with purified StART domain, subjected to UV crosslinking and visualized by in-gel fluorescence of the clicked fluorophore. **D** quantification of +UV/-UV ratios normalized to protein loading of liposomes with and without competitor lipids. Welch two sample t-tests were performed with n.s. p > 0.05 and *p < 0.05. **E** Schematic depiction of the principle of the FRET transfer assay. **F** Representative curves of the FRET experiment as performed at least three times. The presence of 3 mol% sphingosine in donor liposomes slows down the transfer of DHE by purified StART domain (light blue *vs.* light pink curves).

To further investigate the relative interaction strength of the StART domain with sphingosine as compared to cholesterol, we performed an *in vitro* competition experiment in which we used liposomes containing pacChol and pacSph together with an excess of natural sphingosine (3-fold excess, thus accounting for 4.5 mol%) or natural cholesterol (20-fold excess, thus accounting for 30 mol% of lipid content). Of note, the maximal concentration of sphingosine used in this experiment was only 4.5 mol% to preserve membrane integrity ^28^. We observed that the modest excess of natural sphingosine decreased StART crosslinking with pacChol (Fig. 3C, quantified in D). On the other hand, a much larger excess of cholesterol could not diminish the crosslinking intensity with pacSph, which indicates a stronger interaction between pacSph and the StART domain of STARD3 in our assay.

To follow the effect of sphingosine on cholesterol transport with a more direct method, we used a FRET-based assay with liposomes containing the fluorescent sterol analogue dehydroergosterol (DHE). The inherent fluorescence of DHE provides a convenient way to monitor its transport directly. In our setup, we created donor liposomes with DHE and fluorescently labeled PE (Dansyl-PE) as a FRET pair while acceptor liposomes were kept non-fluorescent (Fig. 3E). Mixing of the two liposome populations without protein gave a baseline of decreasing FRET signal over time, due to spontaneous exchange of DHE (see Fig. 3F). Upon incubation with the StART-domain, the FRET signal in donor liposomes decreased more rapidly due to protein-mediated DHE transfer. Adding 3 mol% sphingosine to the donor liposomes did not affect the spontaneous DHE exchange but diminished protein-mediated DHE transfer, visualized by a lesser decrease of the FRET signal (Fig. 3F, purple curve). This effect also supports a model in which sphingosine transfer competes with cholesterol transfer.

### Molecular dynamics simulations reveal a head-out orientation of sphingosine during binding and transport

To gain insights into the specific interaction between sphingosine and the binding pocket of the StART domain, we conducted a series of molecular dynamics (MD) simulations. Initially, we placed the StART domain and a sphingosine molecule at varying non-interacting distances within a 10 nm^3^ water box, as depicted in Fig. 4A. The sphingosine molecule displayed a binding affinity throughout the simulations to the protein’s surface (Fig. 4B), inserting itself into the pocket with its hydrophobic tail (Fig. 4B-C, see also Supplementary Video 1) while exposing the hydrophilic head group to the surrounding water environment. During this insertion, sphingosine came into close proximity to S334, thereby supporting the crosslinking results. Notably, to gain access to the cavity, the loop between residues L414 and V342 needed to be in an open conformation to accommodate the tail, as illustrated in Fig. 4A-C. To investigate the membrane - STARD3/sphingosine interaction, we randomly positioned the complex in ten different orientations approximately 2 nm away from a POPC:Cholesterol (70:30) membrane surface (Fig. 4D) and simulated for 1 microsecond. The interaction between STARD3 and the membrane revealed a binding orientation where sphingosine was oriented towards the membrane surface (Fig. 4D, Supplementary Video 2). Furthermore, the protein adjusted its orientation accordingly, aligning itself in a conformation facilitating sphingosine insertion into or release from the membrane as the binding cavity faced the membrane surface (Fig. 4F, Supplementary Video 3). The free energy profile (Fig. 4F) showed that the final protein-membrane orientation, with sphingosine partially inserted into the membrane surface, exhibited a highly favorable energetic state with a free energy value of −9 k_b_T ± 0.5.

**Fig 4.**
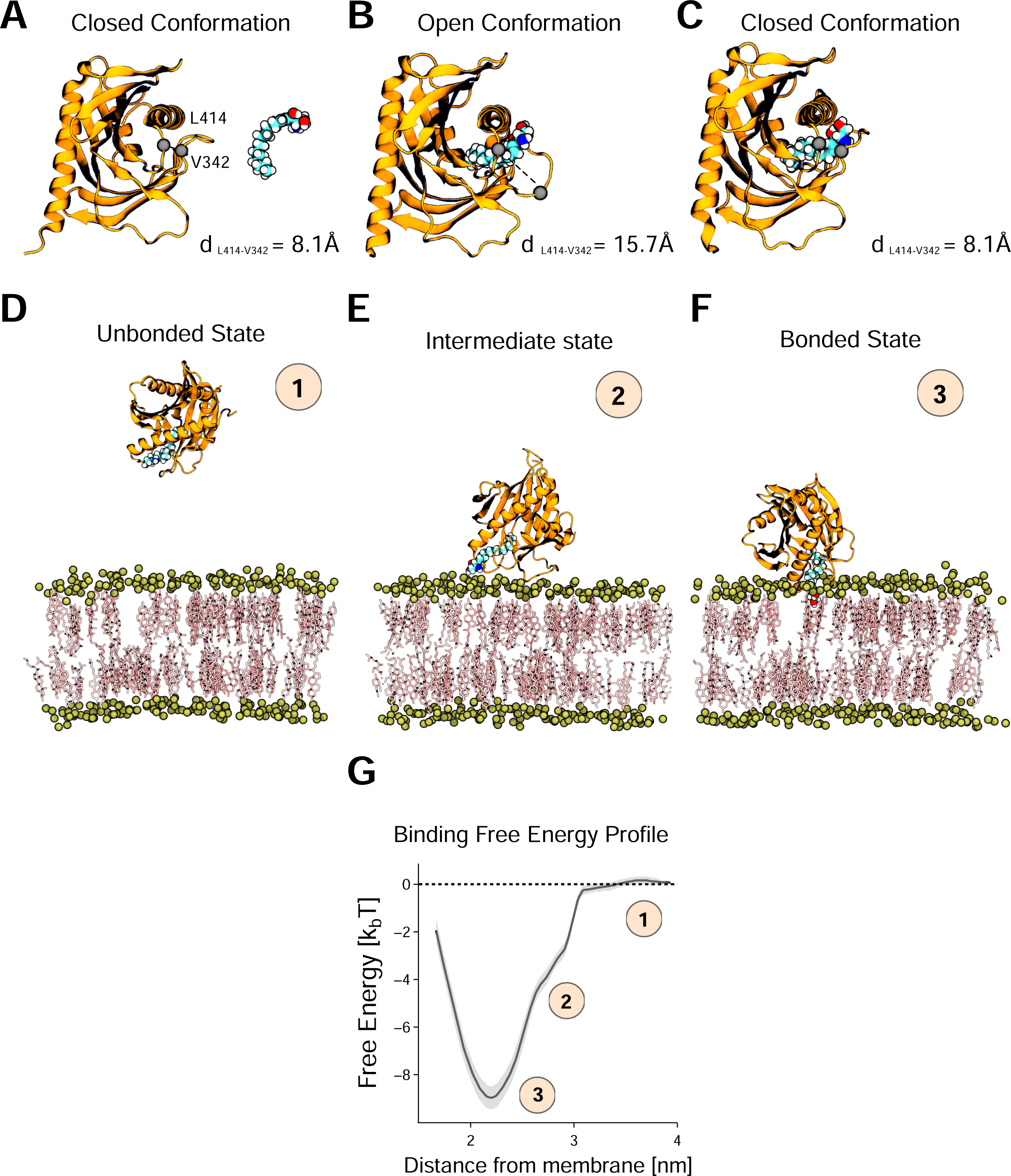
**A-C** Unbiased molecular dynamics simulations snapshots showcasing the spontaneous interaction between sphingosine and STARD3. The simulations were conducted by placing sphingosine at ten different distances from the protein in a non-interactive configuration. As the simulations progressed, sphingosine bound to STARD3 and eventually entered the STARD3 cavity, primarily through its hydrophobic tail. **D-F** Snapshots obtained from unbiased molecular dynamics simulations, highlighting the spontaneous interaction between STARD3 and a model membrane composed of POPC and cholesterol (70:30). In this case, the STARD3-sphingosine complex was positioned at ten different random orientations, initially placed at a non-interactive distance from the membrane surface. The specific orientation of the complex facilitated the interaction of sphingosine with the membrane surface, enabling its insertion or release into or from the membrane as the cavity faced the membrane surface. **G** Free energy profile of the STARD3-sphingosine complex/membrane binding interaction. The profile demonstrates a spontaneous stable (−9 kbT ± 0.5) binding occurring at a distance of approximately 2 nm, measured between the center of mass of the protein and the phosphate atoms of the interacting leaflet in the membrane.

In conclusion, our extensive MD simulations provide valuable insights into the binding mode of sphingosine within the StART domain and its behavior toward a biological membrane. The simulations demonstrated the ability of sphingosine to bind to the protein’s surface and penetrate the pocket with the hydrophobic tail inserted and the hydrophilic head group exposed to the water environment. Moreover, the complex exhibited a preferred orientation towards the membrane surface, accompanied by the protein’s conformational adjustments for potential membrane insertion or release.

## Discussion

Sphingolipids encompass many bioactive lipids which are connected by a delicately balanced metabolic network. Changes to this network can have far-reaching ramifications. For instance, increase in levels of direct sphingosine metabolites, sphingosine-1-phosphate and ceramide, may push the cell towards proliferation or apoptosis, respectively. ^29^ Pathological defects in the sphingolipid metabolic network result in severe diseases, many of which have lysosomal phenotypes, yet our understanding of the molecular architecture of sphingolipid recycling remains incomplete, mainly due to a lack of methods to follow this lipid directly. Here, we show data supporting a model in which the sterol transporter STARD3 binds sphingosine within its StART domain and delivers it to the ER for reuse in the sphingolipid biosynthetic pathway. These findings were made possible by development of technology that facilitated visualization and detection of sphingosine via photocrosslinking. In addition, the ability to photorelease lipid probes in lysosomes of intact cells in a defined manner, gave the necessary spatio-temporal precision to investigate lysosome-to-ER transfer. In this manner, we could show that lysosomally-released sphingosine was incorporated into SM to a higher extent when STARD3 was overexpressed. Interestingly, lipid mass spectrometry did not show changes in SM levels. Given that SM is a highly abundant sphingolipid, particularly at the plasma membrane, we argue that changes in STARD3-dependent SM biosynthesis cannot be resolved when analyzing whole cells in a steady state. For such transport phenomena, out-of-equilibrium approaches are necessary.

To our knowledge, this is the first report of a mammalian interorganellar transfer protein involved in sphingosine trafficking. Protein transporters for ceramides and sphingosine-1-phosphate have been and keep being identified ^30,31^, yet sphingosine transfer proteins remained mysterious. Studies in yeast have recently shown the specific requirement of an ER-vacuole tethering protein, Mdm1/Snx13, in the turnover of vacuolar sphingolipids ^32,33^, but whether this protein is a direct sphingosine transporter remains to be elucidated. It is also a topic of debate whether sphingosine would spontaneously diffuse between organelles ^34^, particularly at sites where two membranes are in close apposition. In our hands, overexpression of a STARD3 construct with defective lipid binding cavity, but functioning tethering motifs did not promote synthesis of complex sphingolipids. Along the same lines, artificial extension of contact sites between lysosomes and ER by expressing a truncated ORP1L construct as a tether also did not fuel lyso-pacSph into the sphingolipid biosynthesis pathway. We therefore argue that, while passive diffusion from lysosomes to ER may occur, transfer of sphingosine is enhanced by the actions of a dedicated transporter such as STARD3.

The identification of the sterol transfer protein STARD3 as a sphingosine transporter follows our previous report which demonstrated that the lysosomal membrane protein NPC1 is capable to export sphingosine as well as sterols through its intramolecular tunnel domain towards the limiting membrane of lysosomes. This would imply that NPC1 acts upstream of STARD3. The subcellular localization of STARD3 was confirmed with late endosomal markers LAMP1, CD63, Rab7, and BMP, which overlap with NPC1-positive endosomes. ^13,35^ Additionally, STARD3 was found to tether lysosomes to mitochondria in cases of NPC1 absence or dysfunction ^8^, implying that contact site formation by STARD3 could be sensitive to lipid cargo provided by NPC1. However, potential co-operation between NPC1 and STARD3 and its implications in sterol and sphingosine transport remain to be experimentally addressed.

Contrary to STARD3 overexpression, the absence of STARD3 resulted only in a mild sphingosine accumulation phenotype, far less pronounced than when lysosomal transporters such as NPC1 ^36^ or Spns1 ^37^ are absent, presuming the export from lysosomal lumen to the lysosomal limiting membrane as the bottle-neck. Considering that sphingosine recycling fulfills a significant portion (50 - 90%) of the demand for long-chain base in the synthesis of cellular sphingolipids ^38^, it is likely that other inter-organellar transporters or additional transport mechanisms such as spontaneous diffusion may share the task of transporting lysosomal sphingosine to target organelles. In addition to redundancy in proteins transporting sphingosine at the lyso-ER interface, it could also be conceivable that sphingosine traffic is rerouted via the PM if the direct lysosome-ER path is blocked, as has been observed for cholesterol. ^39^ Interestingly, the same study found that sphingosine kinases 1 and 2 regulate lysosome-ER membrane contact sites as well as the interorganellar movement of cholesterol, highlighting the importance of lysosomal sphingosine levels in sterol homeostasis and representing another piece of evidence in the ever-growing library for co-regulation of sphingolipid and cholesterol transport and metabolism. ^21,40,41^

The structural features of sphingosine binding inside the sterol-binding lipid transfer cavity of STARD3 were experimentally determined by *in vitro* crosslinking followed by mass spectrometry, which identified a peptide close to the entrance of the lipid cavity. Supporting this, unbiased MD simulations showed that the protein adopted an open conformation by repositioning the loop between residues L414 and V342, the region in which the crosslinked peptide was identified, thereby facilitating sphingosine binding. Additionally, the STARD3 protein was found to come into close proximity with residue S334 during sphingosine insertion, corroborating the experimental crosslinking results. Interestingly, the sphingosine molecule enters the hydrophobic cavity in a “tail-first” orientation. This specific orientation enables the hydrophobic tail of the sphingosine to establish a stable interaction with the internal hydrophobic environment of the protein’s cavity. Concurrently, the hydrophilic head group of sphingosine remains exposed to the surrounding aqueous medium, suggesting a finely tuned mechanism that facilitates effective lipid transfer. MD simulations showed that the sphingosine molecule’s polar head group interacted with the target membrane’s hydrophilic layer upon initial contact with the membrane surface. Following this initial interaction, STARD3 realigned itself, adjusting its orientation so that the binding cavity directly faced the membrane surface. This conformational change was energetically favorable, as evidenced by a free energy value of −9 kbT ± 0.5, implying a stable interaction and potentially facilitating effective lipid transfer between the protein and the membrane.

The tail-first orientation observed in STARD3 for sphingosine’s entry into the hydrophobic cavity is advantageous for lipid release into the target membrane. Recent studies have also confirmed this orientation in other proteins like C1PTP and ORP3 ORD ^42–44^, further supporting the idea that this may be a generalizable mechanism for lipid transfer across various protein systems.

Binding of two ligands to the same protein raises questions of their relative affinities. In absence of dissociation constants for sphingosine and cholesterol, we showed qualitatively that natural sphingosine limited the crosslinking of StART with pacChol. Given that sphingosine also slowed down the transport of dehydroergosterol in a FRET-based lipid transfer assay, we postulate that STARD3 exhibits higher affinity towards sphingosine compared to cholesterol. Higher lipid binding affinity is associated with a slower transport rate, which again points towards the need for other transporters to share the task of transporting lysosomal sphingosine towards the ER. ^45^

The ability to bind multiple ligands, as observed in STARD3, is a relatively common feature of lipid transporters. ^46–49^ Lipid exchangers use this property to transport sterols or phosphatidylserine against their concentration gradient, taking advantage of the hydrolysis of counter-transported lipids such as phosphoinositides. ^48,50–52^ Whether STARD3 also utilizes such a counter-transport mechanism is not clear for now. However, given that sphingosine is rapidly metabolized at the ER by the actions of ceramide synthases and that STARD3 was already shown to deliver cholesterol towards lysosomes ^13^, it is tempting to speculate that some form of counter-transport could take place, further linking sphingolipid and cholesterol metabolic networks.

## Materials and methods

### Protein purification

Sequence of the START domain of STARD3 (STARD3-START, residues 196 – 445) was subcloned into pSFG1 vector (kindly provided by the lab of Ch. Freund, FU Berlin) to generate a His-sfGFP-START fusion protein. A thrombin cleavage site was inserted upstream of the START sequence to enable cleavage between His-sfGFP and STARD3-START domain. Protein was induced in *E. coli*, strain Bl21 (DE3), using induction with cold shock and 0.5 mM IPTG followed by an overnight expression at 18°C, 180 rpm. After expression, bacteria were pelleted and pellets were resuspended in lysis buffer (50 mM Tris pH 7.5, 300 mM NaCl, 20mM imidazole, 10% w/v glycerol) supplemented with protease inhibitor cocktail (Roche). Cells were disrupted using M110 microfluidizer (Microfluidics). Lysate was incubated for 30 min on ice with DNAseI (Roche) and ultracentrifuged at 120 000 g / 4°C / 40 min. His-tagged protein was captured using NiNTA agarose beads (Cube Biotech) and eluted in 10 x 1 mL fractions with elution buffer containing 250 mM imidazole. Fractions with highest protein concentration were pooled together, diluted with thrombin cleavage buffer (20 mM Tris pH 7.5, 150 mM NaCl, 2 mM CaCl_2_) and subjected to cleavage by thrombin (Cytiva) while rotating overnight at 4°C. Cleaved START domain was purified by size exclusion chromatography on a HiLoad 16/60 column (GE Healthcare) equilibrated with 20 mM Tris pH 7.5, 150 mM NaCl, 1 mM DTT. Purified protein fractions were pooled, concentrated, supplemented with 10% glycerol, snap-frozen in liquid nitrogen and stored at −80°C.

### Liposome preparation

Lipids dissolved in chloroform or chloroform-methanol (2:1) were mixed at the desired molar ratio depending on each assay, and the solvent was removed in rotary evaporator (Heidolph) to create a thin lipid film. The films were hydrated with filtered and degassed phosphate buffered saline, (137 mM NaCl, 2.7 mM KCl, 10 mM Na_2_HPO_4_, 1.8 mM KH_2_PO_4_, pH 7.4). Lipid solutions (total lipid concentration = 2 mM) were subjected to 5 freeze-thaw cycles in liquid nitrogen and water (42°C) to generate smaller multilamellar vesicles. These were stored at −20°C. Before use, liposomes were extruded by passing 20x through a 100 nm pore size polycarbonate filter (Whatman - Nuclepore) using a hand extruder (Avanti Polar Lipids). Extruded liposomes were stored at 4°C and used within 1 – 2 days.

### *In vitro* crosslinking assays

Liposomes containing pac-lipids were diluted in PBS until final lipid concentration of 60 µM (with 1.5 µM accessible pac lipid). For *in vitro* crosslinking, most liposomes consisted of 93.5 mol% DOPC with 5 mol% DOPS and 1.5 mol% respective pac-lipid. In competition assays, 1.5 mol% pacChol or 1.5 mol% pacSph was used and 20-fold molar excess of natural cholesterol and 3-fold excess of 18:1 sphingosine was used, respectively (see table below for references of all lipids used in the study). Purified proteins were added until final concentration of 1.5 – 2 µM. Protein-liposome mixtures were incubated in 0.5 mL Eppendorf tubes at 37°C / 30 min / 400 rpm, then crosslinked by exposure to UV light for 10 - 15 minutes at 4°C, using a 100 W mercury lamp. After crosslinking, click reaction was performed by adding 1 µl of freshly prepared click mix into the sample tubes. Final concentration of click reagents in tubes was 80 µM CuSO_4_ (Merck), 3 µM TBTA (Merck), 3 µM Picolyl-Alexa647-Azide (Jena Bioscience) and 80 µM ascorbic acid (Merck). Fresh ascorbic acid solution in water was prepared every time and added to the click mix immediately before the start of click reaction. Reaction was allowed to proceed for 2 hours / 37°C / 400 rpm. Then, samples were concentrated in vacuum evaporator for 30 min / 30 °C and supplemented with 4x Lämmli buffer (250 mM Tris pH 6.8, 9.2% SDS, 40% glycerol, 0.2% bromphenol blue, 100 mM DTT). Samples were boiled for 5 min at 95°C and subjected to SDS-PAGE. Alexa647 and Coomassie images were acquired using the Licor Odyssey imaging system.

Lipids.

Lipids
| Reagent | Source | Identifier |
| --- | --- | --- |
| POPC | Avanti Polar Lipids | Cat# 850457P |
| DOPC | Avanti Polar Lipids | Cat# 850375P |
| DOPS | Avanti Polar Lipids | Cat# 840035P |
| Cholesterol | Merck | Cat# C8667 |
| Sphingosine | Avanti Polar Lipids | Cat# 860490P |
| DHE | Avanti Polar Lipids | Cat# 810253P |
| Dansyl-PE | Avanti Polar Lipids | Cat# 810330C |
| Pac-sphingosine | Avanti Polar Lipids | Cat# 900600P |
| Pac-cholesterol | Merck | Cat# 804657 |
| Pac-PC | Avanti Polar Lipids | Cat# 900408P |
| Pac-ceramide | Avanti Polar Lipids | Cat# 900404P |

### Crosslinking mass spectrometry

Photo-crosslinked samples were acetone-precipitated and processed by in-solution digestion. Briefly, pellets were resuspended in 6M urea/2M thiourea, reduced with 2.5mM dithiothreitol for 30 minutes at room temperature and alkylated with 5mM iodoacetamide for 30 minutes. The combined urea/thiourea concentration was brought to 1.5M by dilution with 50mM ammonium bicarbonate and digestion was carried out with chymotrypsin (1:50 w/w ratio) at 24 °C overnight.

Peptides were recovered from the gel pieces with successive washes in acetonitrile (ACN) and 0.1 trifluoroacetic acid (TFA). The organic phase was then removed by evaporation and the sample was acidified by addition of TFA to 0.1%. Peptides were then cleaned up by C18 StageTips and eluted with 80% acetonitrile, 0.1% TFA, dried and stored at −20 °C until LC-MS acquisition.

Liquid chromatography-mass spectrometry (LC-MS) experiments were carried out on a Fusion Lumos Tribrid Mass Spectrometer (Thermo Fisher Scientific) connected to an Ultimate 3000 UHPLC system (Dionex) operating with XCalibur 4.4 and Tune 3.4. Chromatography was performed with mobile 0.1% (v/v) formic acid as mobile phase A, and 80% (v/v) ACN, 0.1% (v/v) formic acid as mobile phase B. The samples were dissolved in 1.6% ACN (Honeywell Fluka), 0.1% formic acid (Honeywell Fluka) and separated on an Easy-Spray column (C-18, 50 cm, 75 µm internal diameter, 2 µm particle size, 100 Å pore size) running with 300 nl/min flow rate. Approximately 1 µg of peptides were injected per run. The acquisition was carried out with a gradient optimised for the detection of crosslinked species: from 2% B to 11%B in 10 minutes, then to 39%B in 87 min, followed by a ramp to 55% B in 5 minutes and a 15-minute wash at 95%B.

The MS data were acquired in data-dependent mode with a 2.5 second cycle time. Both MS1 and MS2 were acquired in the orbitrap. The following settings were used: MS1 resolution 120,000 with scan range 400-1600 m/z, maximum injection time 50 ms and normalized automatic gain control (AGC) target of 250%. The RF lens was set to 35%. The MS2 was triggered for charge states 3-7 and a single count dynamic exclusion at 60 seconds. Intensity threshold for MS2 triggering was set to 2.5e4. Precursor isolation window was set to 1.4 /m/z and isolation was carried out in the quadrupole. Fragmentation priority was selected with a decision tree strategy ^53^ prioritizing charge states 4-7 over charge state 3. MS2 spectra were acquired with a resolution of 60,000, maximum injection time of 120 ms and normalized AGC of 220%. Fragmentation was carried out with stepped high-energy collision dissociation (HCD) with normalized collision energies of 26,28,30%.

### Mass spectrometry data analysis

Mass spectrometry raw data was searched with MaxQuant (version 2.1.4.0) against the sequence of tagged STARD3 and common contaminants. “Chymotrypsin+” was selected as the protease. Carbamidomethylation of cysteine was set as a fixed modification. Variable modifications included oxidation of methionine, acetylation of protein N-termin. PacSph was defined as a variable modification that can react on any amino acid (composition: H_33_O_2_C_19_N, monoisotopic mass shift: 307.2512). Results were visualized with xiSpec. ^54^

### Data availability

Mass spectrometry data is available at ProteomeXChange PRIDE with the accession number XXX (reviewer code: YYY).

### Generation of cells stably overexpressing STARD3, STARD3 mutants and ORP1L-ΔORD

HeLa 11ht cells stably overexpressing STARD3 and STARD3 mutants were prepared by transfection with a PSF3 vector (a kind gift from Julien Béthune, BZH Heidelberg, DE). The sequence of N-terminally FLAG-tagged STARD3 and C-terminally FLAG-tagged ORP1L-ΔORD were amplified by PCR, introducing EcoRI and NotI restriction sites at the N- and C-terminus, respectively. The PSF3 vector was digested for 1h/37°C using EcoRI and NotI. The amplified PCR product was inserted into the PSF3 vector by ligation with T4 ligase overnight at 16°C.

HeLa 11ht cells, CCL-2, stably expressing the reverse tetracycline-controlled transcription activator rtTA-M2 and containing a locus for Flp-recombinase-mediated cassette exchange ^55,56^ were cultured in Dulbecco’s Modified Eagle’s Medium (Merck) supplemented with 10% FCS (Bio & Sell) and 1% Penicilin/Streptomycin (Merck). Cells were grown in a humidified incubator at 37°C with 5% CO_2_ and regularly tested for mycoplasma contamination. To generate stably overexpressing cell lines, HeLa 11ht cells were grown in a 6-well plate to 70 – 80% confluency and co-transfected with 2 µg of PSF3 vector using polyethylenimine (PEI) in a 1:2 ratio in recovery medium. After 24h, the cells were transferred into a 10 cm dish. After 2 days, the recovery medium was replaced with HeLa 11ht selection medium containing 10% FCS, 1% Penicilin/Streptomycin and 50 mM Ganciclovir. Cells were cultured for one week in selection medium and then seeded into a new 10 cm dish at a very low density. Approximately after one week, colonies of around 20 – 100 cells were grown, picked using sterile cloning discs and expanded to 10 cm dishes. Cells were genetically validated by extracting the genomic DNA (Blood and Tissue Kit, Qiagen), amplifying the stably integrated gene by PCR and sequencing the PCR product.

### Generation of polyclonal CRISPR Cas9 STARD3 knock-out cells

Gene disruption strategy used to produce CRISPR Cas9 knock-out cells was adapted from a protocol by T. Harayama (IPMC, Valbonne, France). ^57^ Empty pX330 and HPRT1-3m13 vectors were a kind gift from T. Harayama. The STARD3 sequence was subcloned into the pX330 vector as described above. HeLa 11ht cells were reverse co-transfected with 495 ng of the pX330 plasmid and 5 ng of HPRT1-3m13 in a 24-plate using Lipofectamine 2000. On the next day, cells were seeded into a 10 cm dish, grown for 4 days and then seeded into a new 10 cm dish with HeLa CRISPR Cas9 selection medium containing 10% FCS, 1% Penicilin/Streptomycin and 6 µg/ml 6-thioguanine. Cells were cultured for one week in the selection medium and afterwards grown in HeLa complete medium. Cells were genetically validated by extracting the genomic DNA, amplifying the exon of interest by PCR and sequencing the PCR product.

### Immunoprecipitation assay

Cells were grown in 10 cm dishes to 90 % confluency and labeled with 2 µM pacSph for 1 h in starvation medium. Thereafter, cells were washed with PBS and 5 mL pre-cooled imaging buffer was added. Lipids were crosslinked to proteins in proximity by irradiating the cells with UV light at 365 nm wavelength for 5 min. Subsequently, cells were harvested by scraping them in PBS and pelleted in a 1.5 mL tube. Cells were resuspended in 100 - 200 µL SDS lysis buffer containing 0.1 % (w/v) SDS and 1 % (v/v) Triton-X-100 in PBS and sonicated for 5 min, followed by a 1 h incubation at 4 °C while rotating. Cells debris was pelleted for 5 min at 14000 rpm at 4 °C and protein-containing supernatant was collected and transferred into a new 1.5 mL tube. Protein concentration was determined using amido black and adjusted to 200 µg protein in 100 µL SDS lysis buffer. 7 µL of click mixture containing 500 µM CuSO_4_, 50 µM TBTA, 500 µM freshly prepared ascorbic acid, and 250 µM picolyl-azide-PEG4-Biotin were added and the solution was pipetted up and down until clear. The click reaction was performed at 37 °C for 1 h while shaking at 500 rpm. Next, to remove all reagents from the click mixture, the proteins were precipitated by adding 400 µL pre-cooled MeOH, 100 µL pre-cooled CHCl_3_ and 300 µL pre-cooled H_2_O, thoroughly vortexed, followed by a centrifugation step at 14000 rpm for 2 min at 4 °C. Two phases were formed, and the protein pellet was located on the actual phase. The upper phase was carefully removed and further 400 µL pre-cooled MeOH was added. The mixture was briefly vortexed and centrifuged again for 3 min at 14000 rpm at 4 °C. The protein pellet was now found on the bottom of the tube and the supernatant was removed. The protein pellet was dissolved in 2 % SDS/PBS by pipetting up and down and incubated for 5 min at 37 °C while shaking. After the pellet was completely dissolved, the precipitation was repeated. This time, after the second centrifuging step, the protein pellet was air-dried for 5 min to evaporate all solvents. Afterwards, the dried protein pellet was resuspended in 200 µL 0.2 % (w/v) SDS/PBS by pipetting up and down, followed by an incubation at 37 °C while shaking until the solution was clear. 20 µL (10 %) were taken as input and the remaining protein was incubated overnight with 20 µL Streptavidin Sepharose beads (Cytiva) previously equilibrated in 0.2 % (w/v) SDS/PBS. On the next day, beads were spun down for 1 min at 1000 rpm and 20 µL of the supernatant was taken as flow-through. Beads were washed twice with 1 mL of IP wash buffer 1 (2 % (w/v) SDS), and twice with IP wash buffer 2 (50 mM HEPES pH 7.4, 1 mM EDTA, 500 mM NaCl, 1 % (v/v) Triton-X-100, 0.1 % (w/v) Na-deoxycholate). Protein was eluted by incubating the cells with 20 µL of elution buffer (10 mM Tris pH 7.4, 2 % (w/v) SDS, 5 % (v/v) β-mercaptoethanol, 2 mM biotin) for 15 min at 98 °C while shaking. Beads were removed by spinning them through a Mobicol “Classic” column with an inserted 10 µm filter (Mo Bi Tec) and flow-through was taken as eluate. Afterwards, 7 µL 4×lämmli buffer was added to input, flow-through and eluate and boiled for further 5 min at 98 °C and western blotting was performed according to standard protocols.

### Microscopy of clickable lipids

Cells were grown on 25 mm glass coverslips placed in a 24-well plate to 60-70 % confluency, labeled with 10 µM lyso-pacSph for 1 h in pre-warmed complete medium and chased overnight in complete medium to allow its pre-localization to lysosomes. The next day, the probe taken up from the cells was uncaged for 90 sec at 405 nm wavelength in pre-warmed imaging buffer. After chasing in imaging buffer at 37 °C, the probe was crosslinked for 5 min in pre-cooled imaging buffer to proteins and other cellular material in close proximity using UV light at 365 nm wavelength. Cells were immediately fixed with MeOH for 20 min at −20 °C. Following fixation, lipids not crosslinked to cellular material were extracted by washing three times with CHCl_3_/MeOH/AcOH (=10:55:0.75 (v/v)) and two times with PBS. Cells were then incubated with 50 µL of click mixture containing 2 mM Cu(I)BF_4_, 0.8 µM AlexaFluor594-picolyl-azide in PBS for 1 h at room temperature in the dark. Cells were then washed two times with PBS and incubated with 50 µL of primary antibody (α-LAMP1 rabbit, Cell Signaling, 1:100 in 1 % BSA, 0.3 % Triton-X-100 in PBS) for 1 h at RT in the dark. Coverslips were washed briefly with PBS and cells were incubated with 50 µL of secondary antibody (α-rabbit conjugated with AlexaFluor488, Cell Signaling, 1:800 in 1 % BSA, 0.3 % Triton-X-100 in PBS) for 30 min at room temperature in the dark. Coverslips were washed briefly with PBS and mounted in 5 µL ProLong Gold Antifade mounting medium (Thermo Fisher Scientific). Cells were imaged at room temperature using a confocal laser scanning microscope (Zeiss LSM800) with a 63× oil objective. Co-localization was analyzed by calculating the Pearson’s correlation coefficient using the Coloc2 tool in the Fiji software (http://fiji.sc/Fiji).

### Thin-layer chromatography of clickable lipids

Cells were grown in a 12-or 6-well plate to 90 % confluency and labeled with different functionalized lipids. For general investigations of cellular sphingolipid metabolism, cells were pulsed for 5 min with 2 µM pacSph and chased for indicated times in starvation medium. For a more precise analysis of Sph export out of the lysosome, a new lyso-pacSph probe was made by Janathan Juarez and Judith Notbohm in the Höglinger laboratory ^21^. This probe was given to cells for 1 h at a 5 µM concentration and chased overnight in complete medium to allow its pre-localization to lysosomes. On the next day, an uncaging step was performed for 90 sec at 405 nm wavelength to activate the probe and then a second chase was done for indicated times to precisely analyze its metabolism after lysosomal export. Subsequently after the chase (pacSph) or the second chase (lyso-pacSph), cells were scraped on ice and pelleted in a 1.5 mL tube at 1500 rpm for 5 min at 4 °C.

Lipids were extracted using a two-phase lipid extraction method. In the first step, cells were resuspended in 300 µL PBS and afterwards 600 µL MeOH and 150 µL CHCl_3_ were added. By centrifuging at 14000 rpm for 5 min, cell debris was pelleted and the supernatant containing the lipids was transferred to a 2 mL tube. Next, 300 µL CHCl_3_ and 600 µL 0.1 % (v/v) acetic acid were added to purify the lipid mixture. After a second centrifugation step at 14000 rpm for 5 min to phases were formed. The aqueous upper phase and remaining protein at the actual phase were taken off and the lipid-containing lower organic phase was transferred in a new 1.5 mL tube. Lipids were dried using a Speed Vac at 30 °C for 20 min. Next, the dried lipids were dissolved in a 30 µL click-mixture containing 4.3 µM 3-azido-7-hydroxycoumarin, 2 mM Cu(I)BF_4_ in EtOH. Click reaction was performed in a speed vac at 45 °C for 20 min. Clicked lipids were dissolved in 15 µL EtOH/acetonitrile (=5:1 (v/v)) and applied on a 10 × 20 cm TLC Silica gel 60 aluminum plate. TLC plates were developed using two different solvents. The plates were placed into a glass chamber containing the first solvent, CHCl_3_/MeOH/H_2_O/AcOH (=65:25:4:1 (v/v)), until the capillary force pushed the solvent front to 5 cm from the bottom. The plates were dried and afterward placed into a second glass chamber containing the second solvent, cyclohexane/ethylacetate (=1:1 (v/v)). After the solvent front was run to the top of the plate, the plate was dried again and lipids containing the fluorescent coumarin group were visualized with a Gel Doc system.

The amount of a certain lipid species was analyzed by calculating the ratio of the lipid species of interest and all lipid species having the fluorescent coumarin group and presented as percent of all lipid species.

### Quantification of ceramide by mass spectrometry

1 × 10^6^ cells were seeded into 10 cm dishes and grown for 2 days. Cells were scraped in PBS and counted using Neubauer Improved cell counting chambers. Cells were pelleted for 5 min at 1500 rpm, supernatant was removed, and cell pellet was snap frozen in liquid nitrogen.

Then 2 mL of cold methanol was added along with the internal standard cocktail (250 pmol of each species dissolved in a final total volume of 10 µL of ethanol:methanol:water 7:2:1). The contents were dispersed using an ultra sonicator at room temperature for 30 s and 1 mL CHCl3 was added. The extract was centrifuged and the supernatant was removed by a Pasteur pipette and transferred to a new tube. The extract was reduced to dryness using a Speed Vac. The dried residue was reconstituted in 0.5 ml of the starting mobile phase solvent for LC-MS/MS analysis, sonicated for ca 15 sec, then centrifuged for 5 min in a tabletop centrifuge before transfer of the clear supernatant to the autoinjector vial for analysis as previously described ^39^. Ceramide was quantified by liquid chromatography electrospray ionization-tandem mass spectrometry (LC-ESI-MS/MS, 5500 QTRAP, ABI) ^39^. Ceramide levels were presented as pmol/mg protein.

### Molecular dynamics simulations

We utilized residues 231-444 from the crystallographic structure of the cholesterol-binding domain of human STARD3 (PDB ID: 5I9J). ^58^ However, the C-terminal residue A445 was absent in this structure, so we employed the GalxyFill software ^59^ through the CHARMM-GUI web server ^60^ to model it. For the C-N termini, we modeled them as charged residues. We derived the Sphingosine 3D structure and parameters from the ceramide 16:0 (CER160) CHARMM-GUI lipid library by removing palmitic acid and appropriately adjusting the partial charge of the amino group. Initially, the protein was subjected to a 1 μs simulation in water with a 0.15M KCl salt concentration. Subsequently, the resulting structure was placed 2 nm away from a randomly oriented sphingosine molecule, and this setup was repeated for ten different random orientations, with each simulation running for 1 μs. At ten random orientations, one protein-sphingosine complex final structure was placed 2nm from a pre-built model membrane surface composed of POPC:CHOL (70:30). This model membrane was built using the CHARMM-GUI membrane builder. ^61^ The ten replicates were solvated with 34,408 water molecules and a 0.15M KCl salt concentration. The system’s charge was neutralized by adjusting the number of K+ or Cl− ions accordingly. Subsequently, each system underwent a 1 μs simulation under NpT conditions. During the production run, we employed the Parrinello-Rahman barostat ^62^ with a semi-isotropic pressure coupling scheme and a time constant of 5.0 ps to maintain constant pressure. The pressure was set to 1.0 bar, and the isothermal compressibility to 4.5 × 10−5 bar−1. Temperature was held at 310 K using the Nose-Hoover thermostat ^63^ with a time constant of 1.0 ps. Electrostatic interactions were calculated using the PME method ^64^, employing a cutoff length of 1.2 nm for both electrostatic and van der Waals interactions. The LINCS algorithm ^65^ constrains hydrogen bonds. The simulations were performed under periodic boundary conditions in all dimensions, utilizing a time step of 2 fs, and coordinates were saved every 100 ps. The GROMACS-2022 software ^66^ was employed for conducting the simulations. The protein, lipids and salt ions were described using the CHARMM36m force field ^67–69^ while the TIP3 model ^70^ was utilized for water molecules. Visual molecular dynamics (VMD) software ^71^ was used for rendering snapshots, and movies.

### Free energy calculation

We performed biased atomistic molecular dynamics simulations utilizing the umbrella sampling protocol to determine the potential of mean force (PMF) for binding STARD3/sphingosine complex on a model membrane surface ^72,73^. The initial configurations for each umbrella window were obtained directly from previous unbiased MD simulations. The reaction coordinate was the center of mass distance between STARD3 and the phosphate atoms of one leaflet in the membrane. A total of 45 umbrella windows, spaced 0.1 nm apart, were generated and simulated with a harmonic restraint force constant of 2,000 kJ/mol/nm^2^ for 200 ns each. The initial 50 ns of each simulation were considered an equilibration phase and subsequently excluded from the actual free energy calculation. The free energy profiles were reconstructed using the weighted histogram analysis method ^74^, which allowed us to estimate the PMF. To assess the statistical error, we performed 200 bootstrap analyses.

## Acknowledgements

This work was supported by the Deutsche Forschungsgemeinschaft (SFB/TRR 186, project A1 and A19; WN and DH, DFG JA 3315/1-1; DJ, DFG LO 2821/1-1; FL) and the National Institutes of Health Grant R01GM043880 (S.S.). The authors gratefully acknowledge the data storage service SDS@hd supported by the Ministry of Science, Research, and the Arts Baden-Württemberg (MWK), the German Research Foundation (DFG) through grant INST 35/1314-1 FUGG and INST 35/1503-1 FUGG. We acknowledge the computing resources provided by the CSC – IT Center for Science Ltd. (Espoo, Finland). The authors acknowledge the Virginia Commonwealth University Lipidomics/Metabolomics, which is supported in part by funding from the NIH-NCI Cancer Center Support Grant P30 CA016059. We thank Jutta Worsch for technical assistance.

## Supplementary Figures

**Supplementary Figure 1:**
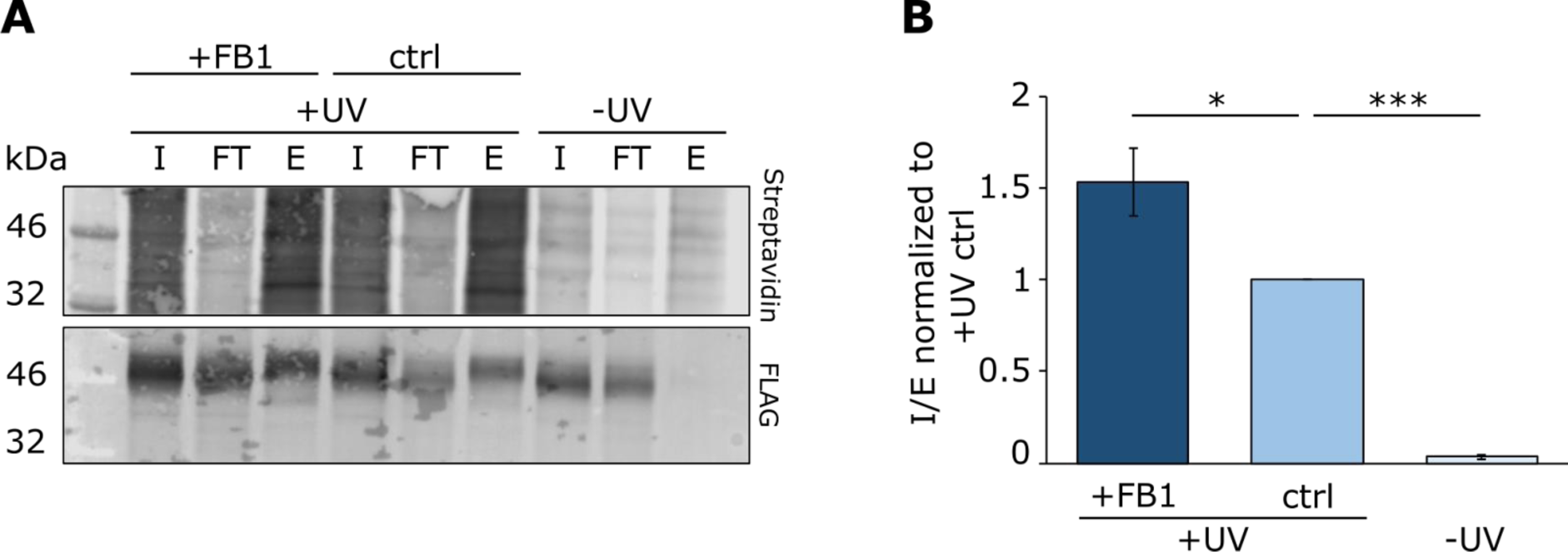
Immunoprecipitation of STARD3 and pacSph in living SGPL1 KO cells. **A** HeLa SGPL1 KO cells were transiently transfected with a N-terminal tagged STARD3-FLAG construct, and +FB1 condition was treated with 50 µM FB1 overnight and additionally 100 µM FB1 1h before the experiment. Cells were labeled with 2 µM pacSph for 1 h, and lipid-protein complexes were crosslinked for 5 min with UV light (365 nm). Crosslinked protein-lipid complexes were conjugated with biotin-azide and bound to streptavidin-coupled beads. Eluted protein-lipid complexes were analyzed by western blotting using Streptavidin and FLAG-specific antibodies. **B** pacSph-STARD3 complexes were quantified by calculating the ratio of I to E normalized to +UV ctrl condition

**Supplementary Figure 2:**
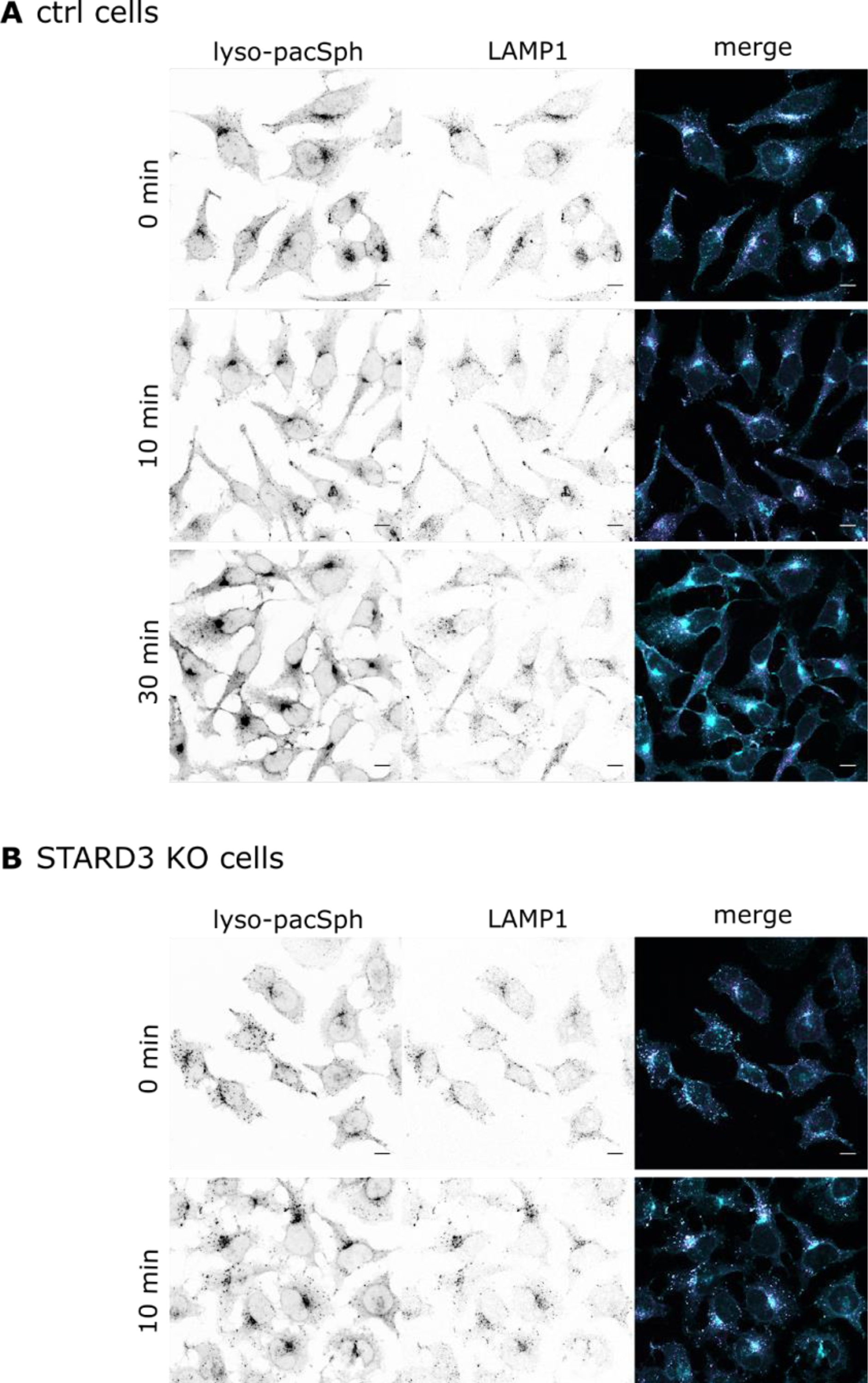
Confocal microscopy images of **A** ctrl and **B** STARD3 KO cells labeled with 10 µM lyso-pacSph. Scale bars, 10 µm. Lyso-pacSph was uncaged by UV light (405 nm) and crosslinked (365 nm) to proteins in close proximity 0 - 30 min post uncaging. Lyso-pacSph metabolites were conjugated to an Alexa488 fluorophore, and lysosomes were visualized using an immunostaining against LAMP1. Microscopy images show the subcellular lyso-pacSph or LAMP1 distribution, while merge images show lyso-pacSph (cyan) and LAMP1 (magenta) co-localization.

**Supplementary Figure 3:**
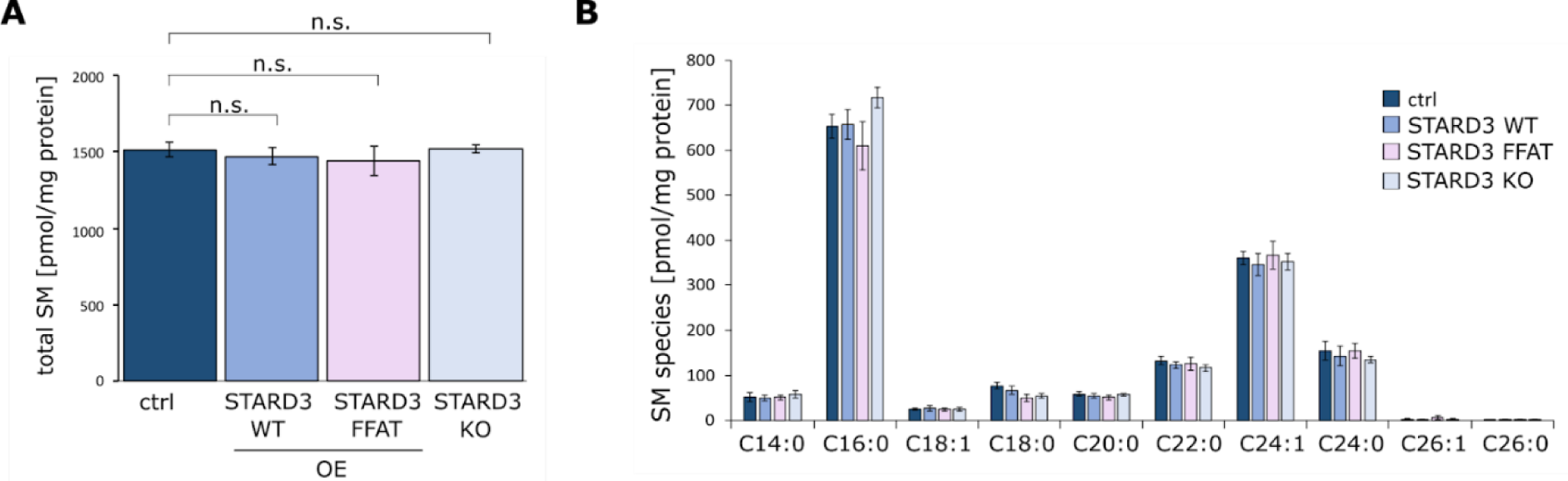
Lipid mass spectrometry results of **A** total SM levels and **B** single SM species performed using total cell lipid extracts of ctrl, STARD3 OE, STARD3 FFAT mutant OE, and STARD3 KO cells. Welch two sample t-tests were performed with n.s. p > 0.05, **p < 0.01 and ***p < 0.001.

## References

1. Lawrence, R.E., and Zoncu, R. (2019). The lysosome as a cellular centre for signalling, metabolism and quality control. Nat. Cell Biol. 21, 133–142. 10.1038/s41556-018-0244-7.

2. Platt, F.M., d’Azzo, A., Davidson, B.L., Neufeld, E.F., and Tifft, C.J. (2018). Lysosomal storage diseases. Nat Rev Dis Primers 4, 27. 10.1038/s41572-018-0025-4.

3. Grassi, S., Chiricozzi, E., Mauri, L., Sonnino, S., and Prinetti, A. (2019). Sphingolipids and neuronal degeneration in lysosomal storage disorders. J. Neurochem. 148, 600–611. 10.1111/jnc.14540.

4. Di Pardo, A., and Maglione, V. (2018). Sphingolipid Metabolism: A New Therapeutic Opportunity for Brain Degenerative Disorders. Front Neurosci 12, 249. 10.3389/fnins.2018.00249.

5. Davidson, S.M., and Vander Heiden, M.G. (2017). Critical Functions of the Lysosome in Cancer Biology. Annu Rev Pharmacol Toxicol 57, 481–507. 10.1146/annurev-pharmtox-010715-103101.

6. Ge, W., Li, D., Gao, Y., and Cao, X. (2015). The Roles of Lysosomes in Inflammation and Autoimmune Diseases. Int Rev Immunol 34, 415–431. 10.3109/08830185.2014.936587.

7. Schoop, V., Martello, A., Eden, E.R., and Höglinger, D. (2021). Cellular cholesterol and how to find it. Biochim Biophys Acta Mol Cell Biol Lipids 1866, 158989. 10.1016/j.bbalip.2021.158989.

8. Höglinger, D., Burgoyne, T., Sanchez-Heras, E., Hartwig, P., Colaco, A., Newton, J., Futter, C.E., Spiegel, S., Platt, F.M., and Eden, E.R. (2019). NPC1 regulates ER contacts with endocytic organelles to mediate cholesterol egress. Nat Commun 10, 4276. 10.1038/s41467-019-12152-2.

9. Qian, H., Wu, X., Du, X., Yao, X., Zhao, X., Lee, J., Yang, H., and Yan, N. (2020). Structural Basis of Low-pH-Dependent Lysosomal Cholesterol Egress by NPC1 and NPC2. Cell 182, 98–111.e18. 10.1016/j.cell.2020.05.020.

10. Heybrock, S., Kanerva, K., Meng, Y., Ing, C., Liang, A., Xiong, Z.-J., Weng, X., Ah Kim, Y., Collins, R., Trimble, W., et al. (2019). Lysosomal integral membrane protein-2 (LIMP-2/SCARB2) is involved in lysosomal cholesterol export. Nat Commun 10, 3521. 10.1038/s41467-019-11425-0.

11. Du, X., Kumar, J., Ferguson, C., Schulz, T.A., Ong, Y.S., Hong, W., Prinz, W.A., Parton, R.G., Brown, A.J., and Yang, H. (2011). A role for oxysterol-binding protein-related protein 5 in endosomal cholesterol trafficking. J Cell Biol 192, 121–135. 10.1083/jcb.201004142.

12. Zhao, K., and Ridgway, N.D. (2017). Oxysterol-Binding Protein-Related Protein 1L Regulates Cholesterol Egress from the Endo-Lysosomal System. Cell Rep 19, 1807–1818. 10.1016/j.celrep.2017.05.028.

13. Wilhelm, L.P., Wendling, C., Védie, B., Kobayashi, T., Chenard, M.-P., Tomasetto, C., Drin, G., and Alpy, F. (2017). STARD3 mediates endoplasmic reticulum-to-endosome cholesterol transport at membrane contact sites. EMBO J 36, 1412–1433. 10.15252/embj.201695917.

14. Lim, C.-Y., Davis, O.B., Shin, H.R., Zhang, J., Berdan, C.A., Jiang, X., Counihan, J.L., Ory, D.S., Nomura, D.K., and Zoncu, R. (2019). ER-lysosome contacts enable cholesterol sensing by mTORC1 and drive aberrant growth signalling in Niemann-Pick type C. Nat Cell Biol 21, 1206–1218. 10.1038/s41556-019-0391-5.

15. Radulovic, M., Wenzel, E.M., Gilani, S., Holland, L.K., Lystad, A.H., Phuyal, S., Olkkonen, V.M., Brech, A., Jäättelä, M., Maeda, K., et al. (2022). Cholesterol transfer via endoplasmic reticulum contacts mediates lysosome damage repair. EMBO J 41, e112677. 10.15252/embj.2022112677.

16. Kitatani, K., Idkowiak-Baldys, J., and Hannun, Y.A. (2008). The sphingolipid salvage pathway in ceramide metabolism and signaling. Cell. Signal. 20, 1010–1018. 10.1016/j.cellsig.2007.12.006.

17. Maceyka, M., and Spiegel, S. (2014). Sphingolipid metabolites in inflammatory disease. Nature 510, 58–67. 10.1038/nature13475.

18. Quinville, B.M., Deschenes, N.M., Ryckman, A.E., and Walia, J.S. (2021). A Comprehensive Review: Sphingolipid Metabolism and Implications of Disruption in Sphingolipid Homeostasis. IJMS 22, 5793. 10.3390/ijms22115793.

19. Höglinger, D., Nadler, A., Haberkant, P., Kirkpatrick, J., Schifferer, M., Stein, F., Hauke, S., Porter, F.D., and Schultz, C. (2017). Trifunctional lipid probes for comprehensive studies of single lipid species in living cells. Proc. Natl. Acad. Sci. U.S.A. 114, 1566–1571. 10.1073/pnas.1611096114.

20. Höglinger, D. (2019). Bi-and Trifunctional Lipids for Visualization of Sphingolipid Dynamics within the Cell. Methods Mol. Biol. 1949, 95–103. 10.1007/978-1-4939-9136-5_8.

21. Altuzar, J., Notbohm, J., Stein, F., Haberkant, P., Hempelmann, P., Heybrock, S., Worsch, J., Saftig, P., and Höglinger, D. (2023). Lysosome-targeted multifunctional lipid probes reveal the sterol transporter NPC1 as a sphingosine interactor. Proc Natl Acad Sci U S A 120, e2213886120. 10.1073/pnas.2213886120.

22. Tomasetto, C., Régnier, C., Moog-Lutz, C., Mattei, M.G., Chenard, M.P., Lidereau, R., Basset, P., and Rio, M.C. (1995). Identification of four novel human genes amplified and overexpressed in breast carcinoma and localized to the q11-q21.3 region of chromosome 17. Genomics 28, 367–376. 10.1006/geno.1995.1163.

23. Alpy, F., Rousseau, A., Schwab, Y., Legueux, F., Stoll, I., Wendling, C., Spiegelhalter, C., Kessler, P., Mathelin, C., Rio, M.-C., et al. (2013). STARD3 or STARD3NL and VAP form a novel molecular tether between late endosomes and the ER. J Cell Sci 126, 5500–5512. 10.1242/jcs.139295.

24. Di Mattia, T., Wilhelm, L.P., Ikhlef, S., Wendling, C., Spehner, D., Nominé, Y., Giordano, F., Mathelin, C., Drin, G., Tomasetto, C., et al. (2018). Identification of MOSPD2, a novel scaffold for endoplasmic reticulum membrane contact sites. EMBO Rep 19. 10.15252/embr.201745453.

25. Gerl, M.J., Bittl, V., Kirchner, S., Sachsenheimer, T., Brunner, H.L., Lüchtenborg, C., Özbalci, C., Wiedemann, H., Wegehingel, S., Nickel, W., et al. (2016). Sphingosine-1-Phosphate Lyase Deficient Cells as a Tool to Study Protein Lipid Interactions. PLoS One 11, e0153009. 10.1371/journal.pone.0153009.

26. Rocha, N., Kuijl, C., van der Kant, R., Janssen, L., Houben, D., Janssen, H., Zwart, W., and Neefjes, J. (2009). Cholesterol sensor ORP1L contacts the ER protein VAP to control Rab7-RILP-p150 Glued and late endosome positioning. J Cell Biol 185, 1209–1225. 10.1083/jcb.200811005.

27. Geiger, T., Wehner, A., Schaab, C., Cox, J., and Mann, M. (2012). Comparative proteomic analysis of eleven common cell lines reveals ubiquitous but varying expression of most proteins. Mol Cell Proteomics 11, M111.014050. 10.1074/mcp.M111.014050.

28. Contreras, F.-X., Sot, J., Alonso, A., and Goñi, F.M. (2006). Sphingosine Increases the Permeability of Model and Cell Membranes. Biophysical Journal 90, 4085–4092. 10.1529/biophysj.105.076471.

29. Newton, J., Lima, S., Maceyka, M., and Spiegel, S. (2015). Revisiting the sphingolipid rheostat: Evolving concepts in cancer therapy. Exp Cell Res 333, 195–200. 10.1016/j.yexcr.2015.02.025.

30. Limar, S., Körner, C., Martínez-Montañés, F., Stancheva, V.G., Wolf, V.N., Walter, S., Miller, E.A., Ejsing, C.S., Galassi, V.V., and Fröhlich, F. (2023). Yeast Svf1 binds ceramides and contributes to sphingolipid metabolism at the ER cis-Golgi interface. J Cell Biol 222, e202109162. 10.1083/jcb.202109162.

31. Hanada, K., Kumagai, K., Yasuda, S., Miura, Y., Kawano, M., Fukasawa, M., and Nishijima, M. (2003). Molecular machinery for non-vesicular trafficking of ceramide. Nature 426, 803–809. 10.1038/nature02188.

32. Girik, V., Feng, S., Hariri, H., Henne, W.M., and Riezman, H. (2022). Vacuole-Specific Lipid Release for Tracking Intracellular Lipid Metabolism and Transport in Saccharomyces cerevisiae. ACS Chem Biol 17, 1485–1494. 10.1021/acschembio.2c00120.

33. Henne, W.M., Zhu, L., Balogi, Z., Stefan, C., Pleiss, J.A., and Emr, S.D. (2015). Mdm1/Snx13 is a novel ER-endolysosomal interorganelle tethering protein. J Cell Biol 210, 541–551. 10.1083/jcb.201503088.

34. Somerharju, P. (2015). Is Spontaneous Translocation of Polar Lipids Between Cellular Organelles Negligible? Lipid Insights 8, 87–93. 10.4137/LPI.S31616.

35. Voilquin, L., Lodi, M., Di Mattia, T., Chenard, M.-P., Mathelin, C., Alpy, F., and Tomasetto, C. (2019). STARD3: A Swiss Army Knife for Intracellular Cholesterol Transport. Contact 2, 251525641985673. 10.1177/2515256419856730.

36. Lloyd-Evans, E., Morgan, A.J., He, X., Smith, D.A., Elliot-Smith, E., Sillence, D.J., Churchill, G.C., Schuchman, E.H., Galione, A., and Platt, F.M. (2008). Niemann-Pick disease type C1 is a sphingosine storage disease that causes deregulation of lysosomal calcium. Nat. Med. 14, 1247–1255. 10.1038/nm.1876.

37. Ha, H.T.T., Nguyen, X.T.A., Vo, L.K., Leong, N.C.P., Liu, S., Nguyen, D.T., Lim, P.Y., Wu, Y.J., Nguyen, T.Q., Oh, J., et al. (2022). SPNS1 is required for the transport of lysosphingolipids and lysoglycerophospholipids from lysosomes (Biochemistry) 10.1101/2022.12.14.520377.

38. Gillard, B.K., Clement, R.G., and Marcus, D.M. (1998). Variations among cell lines in the synthesis of sphingolipids in de novo and recycling pathways. Glycobiology 8, 885–890. 10.1093/glycob/8.9.885.

39. Palladino, E.N.D., Bernas, T., Green, C.D., Weigel, C., Singh, S.K., Senkal, C.E., Martello, A., Kennelly, J.P., Bieberich, E., Tontonoz, P., et al. (2022). Sphingosine kinases regulate ER contacts with late endocytic organelles and cholesterol trafficking. Proc Natl Acad Sci U S A 119, e2204396119. 10.1073/pnas.2204396119.

40. Perry, R.J., and Ridgway, N.D. (2006). Oxysterol-binding protein and vesicle-associated membrane protein-associated protein are required for sterol-dependent activation of the ceramide transport protein. Mol Biol Cell 17, 2604–2616. 10.1091/mbc.e06-01-0060.

41. Guan, X.L., Souza, C.M., Pichler, H., Dewhurst, G., Schaad, O., Kajiwara, K., Wakabayashi, H., Ivanova, T., Castillon, G.A., Piccolis, M., et al. (2009). Functional interactions between sphingolipids and sterols in biological membranes regulating cell physiology. Mol Biol Cell 20, 2083–2095. 10.1091/mbc.e08-11-1126.

42. Rogers, J.R., and Geissler, P.L. (2023). Ceramide-1-phosphate transfer protein enhances lipid transport by disrupting hydrophobic lipid–membrane contacts. PLoS Comput Biol 19, e1010992. 10.1371/journal.pcbi.1010992.

43. Simanshu, D.K., Kamlekar, R.K., Wijesinghe, D.S., Zou, X., Zhai, X., Mishra, S.K., Molotkovsky, J.G., Malinina, L., Hinchcliffe, E.H., Chalfant, C.E., et al. (2013). Non-vesicular trafficking by a ceramide-1-phosphate transfer protein regulates eicosanoids. Nature 500, 463–467. 10.1038/nature12332.

44. Tong, J., Tan, L., and Im, Y.J. (2021). Structure of human ORP3 ORD reveals conservation of a key function and ligand specificity in OSBP-related proteins. PLoS ONE 16, e0248781. 10.1371/journal.pone.0248781.

45. Ikhlef, S., Lipp, N.-F., Delfosse, V., Fuggetta, N., Bourguet, W., Magdeleine, M., and Drin, G. (2021). Functional analyses of phosphatidylserine/PI(4)P exchangers with diverse lipid species and membrane contexts reveal unanticipated rules on lipid transfer. BMC Biol 19, 248. 10.1186/s12915-021-01183-1.

46. Chiapparino, A., Maeda, K., Turei, D., Saez-Rodriguez, J., and Gavin, A.-C. (2016). The orchestra of lipid-transfer proteins at the crossroads between metabolism and signaling. Prog. Lipid Res. 61, 30–39. 10.1016/j.plipres.2015.10.004.

47. Raghu, P., Basak, B., and Krishnan, H. (2021). Emerging perspectives on multidomain phosphatidylinositol transfer proteins. Biochimica et Biophysica Acta (BBA) - Molecular and Cell Biology of Lipids 1866, 158984. 10.1016/j.bbalip.2021.158984.

48. Mesmin, B., Bigay, J., Moser von Filseck, J., Lacas-Gervais, S., Drin, G., and Antonny, B. (2013). A four-step cycle driven by PI(4)P hydrolysis directs sterol/PI(4)P exchange by the ER-Golgi tether OSBP. Cell 155, 830–843. 10.1016/j.cell.2013.09.056.

49. Bockelmann, S., Mina, J.G.M., Korneev, S., Hassan, D.G., Müller, D., Hilderink, A., Vlieg, H.C., Raijmakers, R., Heck, A.J.R., Haberkant, P., et al. (2018). A search for ceramide binding proteins using bifunctional lipid analogs yields CERT-related protein StarD7. J Lipid Res 59, 515–530. 10.1194/jlr.M082354.

50. de Saint-Jean, M., Delfosse, V., Douguet, D., Chicanne, G., Payrastre, B., Bourguet, W., Antonny, B., and Drin, G. (2011). Osh4p exchanges sterols for phosphatidylinositol 4-phosphate between lipid bilayers. J Cell Biol 195, 965–978. 10.1083/jcb.201104062.

51. Moser von Filseck, J., Opi, A., Delfosse, V., Vanni, S., Jackson, C.L., Bourguet, W., and Drin, G. (2015). Phosphatidylserine transport by ORP/Osh proteins is driven by phosphatidylinositol 4-phosphate. Science 349, 432–436. 10.1126/science.aab1346.

52. Chung, J., Torta, F., Masai, K., Lucast, L., Czapla, H., Tanner, L.B., Narayanaswamy, P., Wenk, M.R., Nakatsu, F., and De Camilli, P. (2015). PI4P/phosphatidylserine countertransport at ORP5-and ORP8-mediated ER-plasma membrane contacts. Science 349, 428–432. 10.1126/science.aab1370.

53. Kolbowski, L., Mendes, M.L., and Rappsilber, J. (2017). Optimizing the Parameters Governing the Fragmentation of Cross-Linked Peptides in a Tribrid Mass Spectrometer. Anal Chem 89, 5311–5318. 10.1021/acs.analchem.6b04935.

54. Kolbowski, L., Combe, C., and Rappsilber, J. (2018). xiSPEC: web-based visualization, analysis and sharing of proteomics data. Nucleic Acids Research 46, W473–W478. 10.1093/nar/gky353.

55. Schopp, I.M., Amaya Ramirez, C.C., Debeljak, J., Kreibich, E., Skribbe, M., Wild, K., and Béthune, J. (2017). Split-BioID a conditional proteomics approach to monitor the composition of spatiotemporally defined protein complexes. Nat Commun 8, 15690. 10.1038/ncomms15690.

56. Weidenfeld, I., Gossen, M., Löw, R., Kentner, D., Berger, S., Görlich, D., Bartsch, D., Bujard, H., and Schönig, K. (2009). Inducible expression of coding and inhibitory RNAs from retargetable genomic loci. Nucleic Acids Research 37, e50–e50. 10.1093/nar/gkp108.

57. Harayama, T., Hashidate-Yoshida, T., Aguilera-Romero, A., Hamano, F., Morimoto, R., Shimizu, T., and Riezman, H. (2020). Establishment of a highly efficient gene disruption strategy to analyze and manipulate lipid co-regulatory networks (Genetics) 10.1101/2020.11.24.395632.

58. Horvath, M.P., George, E.W., Tran, Q.T., Baumgardner, K., Zharov, G., Lee, S., Sharifzadeh, H., Shihab, S., Mattinson, T., Li, B., et al. (2016). Structure of the lutein-binding domain of human StARD3 at 1.74 Å resolution and model of a complex with lutein. Acta Crystallogr F Struct Biol Commun 72, 609– 618. 10.1107/S2053230X16010694.

59. Coutsias, E.A., Seok, C., Jacobson, M.P., and Dill, K.A. (2004). A kinematic view of loop closure. J Comput Chem 25, 510–528. 10.1002/jcc.10416.

60. Lee, J., Cheng, X., Swails, J.M., Yeom, M.S., Eastman, P.K., Lemkul, J.A., Wei, S., Buckner, J., Jeong, J.C., Qi, Y., et al. (2016). CHARMM-GUI Input Generator for NAMD, GROMACS, AMBER, OpenMM, and CHARMM/OpenMM Simulations Using the CHARMM36 Additive Force Field. J Chem Theory Comput 12, 405–413. 10.1021/acs.jctc.5b00935.

61. Wu, E.L., Cheng, X., Jo, S., Rui, H., Song, K.C., Dávila-Contreras, E.M., Qi, Y., Lee, J., Monje-Galvan, V., Venable, R.M., et al. (2014). CHARMM-GUI Membrane Builder toward realistic biological membrane simulations. J Comput Chem 35, 1997–2004. 10.1002/jcc.23702.

62. Parrinello, M., and Rahman, A. (1981). Polymorphic transitions in single crystals: A new molecular dynamics method. Journal of Applied Physics 52, 7182–7190. 10.1063/1.328693.

63. Hoover, W.G. (1985). Canonical dynamics: Equilibrium phase-space distributions. Phys Rev A Gen Phys 31, 1695–1697. 10.1103/physreva.31.1695.

64. Essmann, U., Perera, L., Berkowitz, M.L., Darden, T., Lee, H., and Pedersen, L.G. (1995). A smooth particle mesh Ewald method. The Journal of Chemical Physics 103, 8577–8593. 10.1063/1.470117.

65. Hess, B., Bekker, H., Berendsen, H.J.C., and Fraaije, J.G.E.M. (1997). LINCS: A linear constraint solver for molecular simulations. J. Comput. Chem. 18, 1463–1472. 10.1002/(SICI)1096-987X(199709)18:12<1463::AID-JCC4>3.0.CO;2-H.

66. Van Der Spoel, D., Lindahl, E., Hess, B., Groenhof, G., Mark, A.E., and Berendsen, H.J.C. (2005). GROMACS: fast, flexible, and free. J Comput Chem 26, 1701–1718. 10.1002/jcc.20291.

67. Pastor, R.W., and Mackerell, A.D. (2011). Development of the CHARMM Force Field for Lipids. J Phys Chem Lett 2, 1526–1532. 10.1021/jz200167q.

68. Huang, J., and MacKerell, A.D. (2013). CHARMM36 all-atom additive protein force field: validation based on comparison to NMR data. J Comput Chem 34, 2135–2145. 10.1002/jcc.23354.

69. Huang, J., Rauscher, S., Nawrocki, G., Ran, T., Feig, M., de Groot, B.L., Grubmüller, H., and MacKerell, A.D. (2017). CHARMM36m: an improved force field for folded and intrinsically disordered proteins. Nat Methods 14, 71–73. 10.1038/nmeth.4067.

70. Jorgensen, W.L., Chandrasekhar, J., Madura, J.D., Impey, R.W., and Klein, M.L. (1983). Comparison of simple potential functions for simulating liquid water. The Journal of Chemical Physics 79, 926–935. 10.1063/1.445869.

71. Humphrey, W., Dalke, A., and Schulten, K. (1996). VMD: visual molecular dynamics. J Mol Graph 14, 33–38, 27–28. 10.1016/0263-7855(96)00018-5.

72. Torrie, G.M., and Valleau, J.P. (1974). Monte Carlo free energy estimates using non-Boltzmann sampling: Application to the sub-critical Lennard-Jones fluid. Chemical Physics Letters 28, 578–581. 10.1016/0009-2614(74)80109-0.

73. Torrie, G.M., and Valleau, J.P. (1977). Nonphysical sampling distributions in Monte Carlo free-energy estimation: Umbrella sampling. Journal of Computational Physics 23, 187–199. 10.1016/0021-9991(77)90121-8.

74. Hub, J.S., De Groot, B.L., and Van Der Spoel, D. (2010). g_wham—A Free Weighted Histogram Analysis Implementation Including Robust Error and Autocorrelation Estimates. J. Chem. Theory Comput. 6, 3713–3720. 10.1021/ct100494z.

